# Prediction of G4 formation in live cells with epigenetic data: a deep learning approach

**DOI:** 10.1101/2023.03.28.534555

**Authors:** Anna Korsakova, Anh Tuân Phan

## Abstract

G-quadruplexes (G4s) are secondary structures abundant in DNA that may play regulatory roles in cells. Despite the ubiquity of the putative G-quadruplex sequences (PQS) in the human genome, only a small fraction forms secondary structures in cells. Folded G4, histone methylation and chromatin accessibility are all parts of the complex *cis* regulatory landscape. We propose an approach for G4 formation prediction in cells that incorporates epigenetic and chromatin accessibility data. The novel approach termed *epiG4NN* efficiently predicts cell-specific G4 formation in live cells based on a local epigenomic snapshot. Our architecture confirms the close relationship between H3K4me3 histone methylation, chromatin accessibility and G4 structure formation. Trained on A549 cell data, *epiG4NN* was then able to predict G4x formation in HEK293T and K562 cell lines. We observe the dependency of model performance with different epigenetic features on the underlying experimental condition of G4 detection. We expect that this approach will contribute to the systematic understanding of correlations between structural and epigenomic feature landscape.

## INTRODUCTION

DNA and RNA are capable of forming multiple conformations of secondary structures, including G-quadruplexes (G4s). G4 structures may be implicated in important biological processes, including replication, transcription (1–3), telomere maintenance (2, 4, 5), RNA processing and translation (6–9). G4-forming sequences are found in the promoters of genes related to cancer, such as *VEGF* (10), *bcl-2* (11), *c-kit* (12, 13), *KRAS* (14). G4 structures are attractive drug targets (15, 16) and it is important to predict their formation in live cells.

The first G4 prediction approaches were based on biophysical knowledge that a DNA motif with 4 runs of at least 3 guanines separated by loops of 1 to 7 nucleotides is likely to fold into a G4, and the first search of such patterns in the human genome resulted in more than 380,000 matches (17, 18). As topological diversity of confirmed G4s was expanded, for example, G4s with long loops (19), G4s with bulges (20) and G4s with missing guanines (21–23), algorithms for search and prediction of G4 evolved to accommodate the diversity of motifs as well. Novel whole-genome searches include inter-molecular G4s formed between the two DNA strands (24), and slightly mismatched sequences (25). Further approaches incorporated contextually enhanced prediction, such as G-runs continuity coupled with loop size (26), and nucleotide content bias (27, 28). Recently, new experimental methods for G4 detection have emerged. A polymerase stop assay Illumina sequencing method was developed that allowed to detect over 525,000 G4s in purified nuclear DNA *in vitro*, or more than 710,000 G4s with a G4-stabilising ligand (29). *In cellulo* G4 detection methods include G4 chromatin immunoprecipitation sequencing (ChIP-seq) methods that use structure-specific antibodies to detect G4 formed in cellular cultures (30–32) and tumor xenografts (33), and G4 detection with cleavage under targets and tagmentation (CUT&Tag) technique (34, 35). The numbers of reported G4s vary greatly between the experimental methods and cell types (Table 1).

**Table 1.** Reported DNA and RNA G4 formed in vitro, in cells and in patient tissue samples.

| GEO ID | Study | Source | Method | Cell, tissue type / experimental conditions | No G4 peaks |
| --- | --- | --- | --- | --- | --- |
| GSE76688 | (31) | HaCaT | BG4 ChIP-seq | epidermal | 19,021 |
| GSE76688 | (31) | NHEK | BG4 ChIP-seq | epidermal | 3,131 |
| GSE99205 | (36) | HaCaT | BG4 ChIP-seq | epidermal | 11,539 |
| GSE107690 | (37) | K562 | BG4 ChIP-seq | blood | 8,955 |
| GSE152216 | (33) | Patient breast cancer samples | qG4 ChIP-seq | breast | 26,000+ |
| GSE133379 | (32) | HEK293T | G4P ChIP-Seq | kidney | 40,790 |
| GSE133379 | (32) | A549 | G4P ChIP-Seq | lung | 123,274 |
| GSE133379 | (32) | H1975 | G4P ChIP-Seq | lung | 152,072 |
| GSE133379 | (32) | HeLa | G4P ChIP-Seq | uterus | 17,787 |
| GSE178668 | (35) | HEK293T | G4-CUT&Tag | kidney | 17,888 |
| GSE178668 | (35) | HEK293T | G4-ChIP-seq | kidney | 9,202 |
| GSE173103 | (38) | HaCaT | G4-CUT&Tag | epidermal | 10,000+ |
| GSE173103 | (38) | HEK293T | G4-CUT&Tag | kidney | 10,000+ |
| GSE181373 | (34) | K562 | G4-CUT&Tag | blood | 21,312 |
| GSE181373 | (34) | U2OS | G4-CUT&Tag | bone | 35,452 |
| GSE63874 | (29) | *in vitro* | G4-seq | *in vitro*, K<sup>+</sup> | 525,908 |
| GSE63874 | (29) | *in vitro* | G4-seq | *in vitro*, K<sup>+</sup>, PDS | 716,311 |
| GSE77282 | (39) | *in vitro* | rG4-seq (RNA) | *in vitro*, K<sup>+</sup> | 3,383 |
| GSE77282 | (39) | *in vitro* | rG4-seq (RNA) | *in vitro*, K<sup>+</sup>, PDS | 11,367 |
| - | This report | *in silico* | Regular expression search | GRCh37/hg19 | 2,105,837 |

Methods for G4 detection in cells imply G4 detection *ex vivo* or *in situ*. The former means that the cells are fixed and chromatin is fragmented before it is enriched with a specific antibody, usually BG4 (30, 31, 36). The latter means that BG4 antibody permeates the plasma and nucleus membranes and tethers Tn5 transposase for tagmentation (35, 38), or another small antibody, G4P, is expressed endogenously for a subsequent ChIP-seq experiment (32). Only a fraction of resulting G4s overlap in the same cell line using two different types methods – about 30-60% (35), or 45% (38). The divergent numbers of G4 formed in different types of cells and detected with different approaches suggest the need for exploration of the biological causes. Predictive models for G4 formation based on these experimental data have been implemented: Quadron (40) and PENGUINN (41) trained on *in vitro* DNA G4 data, DeepG4 (42) with an additional feature of local chromatin accessibility for G4 prediction in cells, and G4RNA screener (43) trained on experimental RNA G4 data. To date, only DeepG4 implemented actual cellular context for G4 formation prediction, however, only the G4s that are formed both *in vitro* and in cells were selected. A significant portion of weaker G4s not folded *in vitro* that might play important regulatory roles in cells were, therefore, excluded from the training data. Excluding weaker G4s might inherently lead to higher accuracies too, as lower experimental scores for these G4s are likely related to finer biological features. Additionally, chromatin accessibility profile was the only feature used. The roles of open chromatin, histone and DNA chemical modifications in G4 formation is being discovered recently (31, 35, 37, 38, 42). Leveraging multiple epigenomic features to predict G4 formation in cells and determining the importance of these features may provide novel insights into G4 formation mechanisms and their regulatory roles.

### G4 existence in the epigenetic context

DNA G-quadruplexes are over-represented in gene promoters and are thought to be involved in gene regulation at the transcription level. More than 40% of gene promoters in the human genome contain G4 forming motifs (44), and their structural properties make them attractive drug targets for diseases involving dysregulation of gene transcription (15). Folded G4s were confirmed to be highly enriched in gene promoters in cells more than in any other region (31), with some studies showing that most folded G4s are, in fact, located in the promoter regions in cells (35). Both G4s formed in cells (31) and gene promoters are associated with open chromatin (45, 46), aiding the accessibility for transcription machinery. Open chromatin was indeed found to contain 85.8% of G4 in HaCaT cells and 97.2% in NHEK cells (31), whereas we found that for A549 cells G4 data (32), only 6.6% of G4s intersected with peaks from ATAC-seq experiment, likely driven by a higher number of G4 peaks reported in A549. It has been hypothesized that G4 formation promotes transcription factor docking by keeping the DNA double helix open (47–49) and allows re-initiation of transcription. Therefore, G4s are likely to be located in accessible chromatin regions due to the local regulatory roles, but the ways G4s are related to accessible chromatin and epigenetic marks are highly complex (Fig. 1). Recently, evidence of high colocalization of other epigenetic marks, such as histone modifications, with G4 formation became available (35). H3K4me3 was found to be the most correlated to G4 formation in HEK293T cells, as measured by G4 ChIP-seq, followed by H3K4me2, H3K4me1 and chromatin accessibility by ATAC-sequencing (35). Additionally, active G4s are present in CpG island (CGIs) regions depleted in cytosine methylation (37, 38) and inhibit methylation of DNA (37). CGI methylation patterns, in turn, mediate binding of specific families of transcription factors that have preference for either methylated or hypomethylated CGI (50), therefore leading to transcriptional regulation via epigenetic modifications. Another mechanism of G4 involvement in cellular processes regulation through epigenetic marking is the G4 involvement in DNA replication. It was demonstrated that G4 formed during DNA replication leads to epigenetic instability due to failure of copying the chromatin repressive marks (51). Additionally, G4 are known to recruit histone modification agents (52, 53) and chromatin remodelling proteins (54). The interplay between folded G4s and epigenetic marks is evident, however, it is still not sufficiently explored for G4 formation prediction.

**Figure 1.**
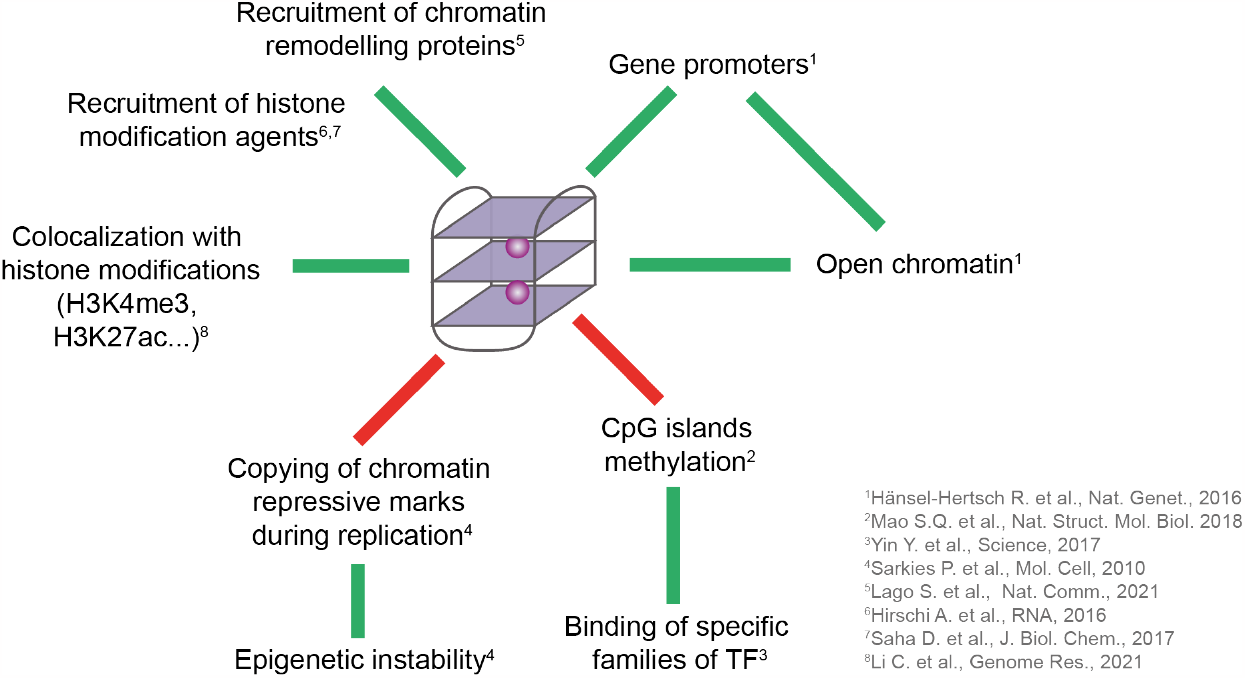
G4 involvement in the epigenetic regulation. G4s are colocalized with gene promoters, open chromatin and certain epigenetic marks (such as H3K4me3), are able to recruit chromatin remodelling proteins and histone modification agents, and prevent CpG islands from cytosine methylation. Additionally, G4 may arrest the copying of the chromatin repressive marks, leading to epigenetic instability.

While the correlations of accessible chromatin and some epigenetic marks hint the relative importance of these features for G4 formation, our goal is to use machine learning to develop a model that can predict G4 formation in cells and rate the importance of these features for G4 formation. Five histone modification marks (H3K4me1, H3K4me3, H3K9me3, H3K27ac, H3K36me3) and chromatin accessibility (ATAC-seq) were selected for training and evaluation. We aim to predict weaker G4s formed in cells along with stable G4s. We focus our predictions on a broad set of putative quadruplex sequences found in the human genome according to the latest motif definitions (see Materials and Methods) and infer the putative quadruplex sequence (PQS) formation probability in cells. The developed approach is designed to transfer the learned features to predict G4 in unseen types of cells based on the underlying sequence and local epigenetic snapshot.

## MATERIAL AND METHODS

### G4 input preparation for epiG4NN model

PQS sequences were found with a regular expression search using *python re* package in the hg19/GRCh37 human genome assembly, retrieved from the UCSC Genome Browser (http://genome.ucsc.edu/). We broadened the existing definitions of G4 and used the following regular expressions: [G_3+_L_1-12_]_3+_G_3+_ – canonical G4 pattern with extended loop length; [GN_0-1_GN_0-1_GL_1-3_]_3+_GN_0-1_GN_0-1_G – bulged G4 pattern with possible G-run breaks; [G_1-2_N_1-2_]_7+_G_1-2_ – irregular G4 pattern, where L is any of {A, T, C, G} and N is any of {A, T, C}. A total number of guanines greater or equal than 12 was required to avoid two-layered G4. The search was performed on both strands. We filtered the redundant and nested G4s by only considering distinct sequences separated by at least one nucleotide. The overlapping sequences were not merged. The first encountered motif from the 5 ‘ end of the overlapping group is considered. A total of 2,105,837 G4 were found for the three types (Supplementary Figure 1). PQS sequences were then padded to 1000 nt and one-hot-encoded for training as follows: A=[1, 0, 0, 0], T=[0, 1, 0, 0], C=[0, 0, 1, 0], G=[0, 0, 0, 1], N=[0, 0, 0, 0]. For each PQS, the respective G4 score was found from the experimental dataset by overlapping the PQS motif with experimental peaks and taking an average of the continuous .*bedgraph* signal. If there is no signal corresponding to the PQS coordinate, the score was set as 0.0. G4 labels for training were obtained from the G4P ChIP-seq study with GEO accession number GSE133379 for A549 and HEK293T cell lines. The upper 5 percentiles of the normalized experimental scores were characterized as “positive” class, and the rest of PQS as “negative” (Supplementary Figure 2). As a result, more than 105,000 PQS were classified as positive for A549 – a slightly more conservative number of peaks as compared to the number of peaks determined in the downstream analyses in the original study (32). A549 cell data were selected for training, and HEK293T for independent evaluation. For HEK293T, upper 2 percentiles were used to match the originally reported number of called peaks (more than 40,000). For additional evaluation and analyses, pre-processed G4 peaks for HeLa (GSE133379), HaCaT (GSE76688) and K562 (GSE107690) were used.

### Chromatin accessibility and epigenetic marks data preparation

Epigenetic information is generally preserved across tissues in species, especially for cell cultures with common progenitor cells (55), therefore, using G4 data and epigenetic data from different studies of the same cell line is possible. We used histone 3 lysine residue 4 methylation and trimethylation, histone 3 lysine 9 and lysine 36 residues trimethylation, histone 3 lysine 27 residue acetylation, and nucleosome availability as epigenetic marks for our experiments. We retrieved H3K4me1, H3K4me3, H3K9me3, H3K27ac, H3K36me3 ChIP-seq datasets and ATAC-seq dataset for A549 cells, and H3K4me3, H3K27ac ChIP-seq for HEK293T from the Encyclopedia of DNA elements (ENCODE) (56) Reference Epigenome project with accession codes ENCFF633KDT, ENCFF021UDY, ENCFF474QYY, ENCFF126BYV, ENCFF053BXF, ENCFF735UWS, and accession codes ENCFF315TAU, ENCFF186KMN, respectively. ATAC-seq signal for HEK293T was retrieved from GEO project with accession code GSM3905877. For K562 cells, we used ENCODE projects with accession codes ENCFF252GZO for ATAC-seq, ENCFF929TPH for H3K4me3, and ENCFF488FYZ for H3K27ac. If the data were originally mapped to hg38, they were lifted over to hg19 using UCSC *liftOver* tool with a chain downloaded from the UCSC genome browser (https://hgdownload.soe.ucsc.edu/goldenPath/hg38/liftOver/). We filtered out regions with no full coverage of any of the features to ensure continuous data availability for every PQS. All epigenetic data were normalized to a range of values [0, 1]. We then created input arrays of 1000-nt epigenetic profiles for each PQS with *bedtools intersect* command-line tool. Genomic tracks were visualized with *pyGenomeTracks* tool(57).

### Model architecture

*epiG4NN* architecture is based on a ResNet (58) – neural network class with residual convolutional layers and dilation. We designed our architecture as sequential convolutional towers of 4 residual blocks each, where each residual block contains batch normalization (x2), rectified linear unit (ReLU) activation (x2), and convolutional layer with 32 kernels and variable kernel size (x2) (Supplementary Figure 3). We constructed a model with only sequence input (*G4NN*) as a baseline for *epiG4NN*. Model inputs are arrays of shape [1000, 4] for *G4NN*, sequence-based model, and arrays of shape [1000, 5] for *epiG4NN*. The first four rows of the input are the one-hot-encoded sequences, and the last row is the normalized epigenetic feature. Output of each residual unit is added to the penultimate layer through a 1D convolution with hidden state of size 32, therefore implementing skip, or residual, connections between units for better convergence and avoidance of vanishing/exploding gradients problem (58). Classification objective for this model is single-class, single label binarized probability of G4 formation. We performed a search for optimal hyperparameter set by training a grid of models with various hyperparameters. The optimal parameters were selected based on the performance of the model on the validation dataset during training: *K* = 3, ***W*** = [11, 11, 15], ***D*** = [1, 4, 10], where K is the number of convolutional towers, *W*_*i*_ is the convolutional filter width in the *i*^*th*^ residual block, *D*_*i*_ is the dilation rate of the *i*^*th*^ residual block.

### Model training and evaluation

Training data from A549 dataset were split into train and test subsets, where the test set contains all the PQS that belong to chromosomes *1, 3, 5, 7, 9*, and train set – all PQS that belong to chromosomes *2, 4, 6, 8, 10-22, X, Y*. The models were trained with batch size of 64 and a constant learning rate of 0.001 with *Adam* stochastic gradient descent optimizer method based on adaptive estimation of lower-order moments (59). Upon initialization, model weights are filled with random numbers. At each training iteration, binary cross entropy loss function is minimized over the target train labels. Output labels are determined via sigmoid function applied to the ultimate logit. Due to the large number of input data samples, chromosome data were input gradually and training on one chromosome from the train set constituted a full iteration. Training then continued with the next chromosome, to a total of 19 iteration steps in one epoch. Optimal hyperparameters were determined from the performance of the sequence-only model on the validation dataset, as determined by accuracy, and applied to all the other models. Most of PQS had zero scores, therefore, the classes are highly imbalanced and are contributing unequally to the metrics and loss, with low probability class (unfolded G4) skewing the metrics. This problem was addressed by implementing balanced class weights for the optimizer: *weight*_*i*_ *= N*_*samples*_*/(N*_*classes*_*·frequency*_*i*_), resulting in weights 0.35 and 35.69 for negative and positive classes. We evaluated the models on withheld test samples using accuracy and area under the receiver operating characteristic curve (AUROC) to determine class separability. AUROC x-axis depicts false positive rate (FPR), and y-axis depicts true positive rate (TPR), defined as *FPR = FP/(FP+TN)* and *TPR = TP/(TP+FN)*. However, these metrics are not optimal for imbalanced classification problems, therefore we additionally evaluate areas under the precision-recall curve (AUPRC) to determine precision and recall of positive G4 samples. Precision (P) and recall (R) are defined as follows: *P = TP/(TP+FP), R = TP/(TP+FN)*, where *TP* is true positives, *FP* is false positives, and *FN* is false negatives. Training and evaluation were performed using TensorFlow (v2.4.0) using python keras API. Respective python scripts can be found at https://github.com/anyakors/epiG4NN.

## RESULTS

### Quadruplex-forming motif definition broadening

Early approaches to defining G4-forming motifs used regular expressions of type [G_3+_L_1-7_]3+G3+ (17, 18, 60) and yielded around 376,000 putative G4s. Many G4s forming *in vitro* still did not adhere to this limited definition (29), including G4s with missing guanines (21–23), G4s with ultra-long loops (19), bulged G4s (20) and G4s with runs of irregular length (61), to name a few. Later search efforts considered variable numbers of guanines in the run (62), bulges and mismatches (“imperfections”) (25, 26), longer loops of up to 12 nucleotides (29), G-rich sequences considering cytosine bias (27), and duplex stem-containing loops (63). As the goal of this work is to define a set of G4 motifs for training that covers a significant part of the *in vivo* formed peaks, we limit ourselves to a few simple definitions. We broadened the existing definitions of G4 and used the following regular expressions: [G_3+_L_1-12_]_3+_G_3+_ – canonical G4 pattern with extended loop length, similarly to (29); [GN_0-1_GN_0-1_GL_1-3_]_3+_GN_0-1_GN_0-1_G – G4 pattern with possible bulges, with restrictions as described in (20); [G_1-2_N_1-2_]_7+_G_1-2_ – irregular G4 pattern alike those studied in (61). These three definitions do not aim to exhaustively cover the G4 motif repertoire, and only selects a representative set of G4 motifs for training. A total of 2,105,837 G-quadruplex motifs were determined after filtering (see Materials and Methods), where 907,845 instances are bulged G4s, 652,908 are irregular G4s, and 545,084 are canonical G4s with extended loop length.

### G4 colocalizes with accessible chromatin and chromatin marks in cells

We interrogated the mean profiles of five epigenetic marks and chromatin accessibility genome-wide at active G4 loci in A549 cells (32) (Fig. 2). Mean normalized signals of H3K4me1, H3K4me3, H3K9me3, H3K27ac, H3K36me3 histone modifications, and ATAC-seq signal were centered and plotted at PQS motifs with G4P ChIP-seq peaks. H3K4me3, H3K27ac and ATAC-seq profiles displayed positive association of epigenetic mark occupancy with the G4 peaks, while H3K4me1, H3K9me3 and H3K36me3 marks demonstrated the opposite trend. H3K4me1, H3K9me3 and H3K36me3 exhibit a dip at G4 sites compared to the genome-wide background level, while H3K4me3, H3K27ac and ATAC-seq demonstrate peaks at G4 sites compared to background level (Fig. 2b). Heatmap analysis (Fig. 2c) of the top 6,000 profiles of each mark revealed that H3K4me3, H3K27ac and open chromatin (ATAC-seq) signals contribute the highest number of informative profiles as well. The profiles were sorted by average intensity and plotted from highest to lowest. H3K4me1, H3K9me3 and H3K36me3 have most of profiles with close-to-zero intensity, while H3K4me3, H3K27ac and ATAC-seq enrich more than six thousand profiles (Fig. 2d).

**Figure 2.**
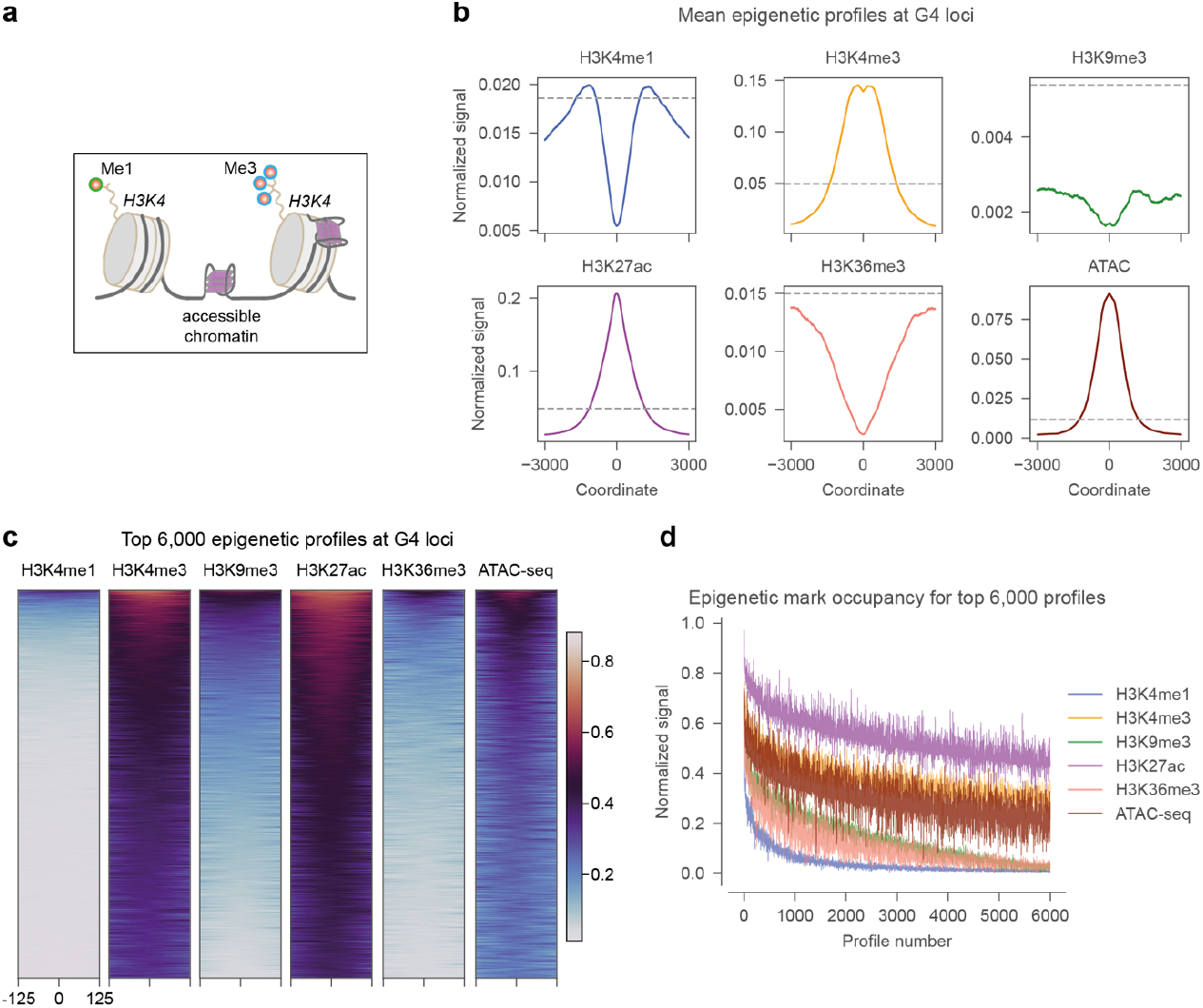
Genome-wide profiles of epigenetic mark occupancy at G4 sites. a) Unlike *in vitro* conditions, in cells G4s are formed in the chromatin context with possible epigenetic chemical modifications, such as histone 3 lysine residue 4 methylation and tri-methylation (H3K4me1, H3K4me3), and others. b) Mean distribution of normalized H3K4me1, H3K4me3, H3K9me3, H3K27ac, H3K36me3 and ATAC-signal (chromatin accessibility) in the 6000 nt vicinity of active G4 loci in A549 cells. The profiles are normalized to have a maximum of 1.0. Mean genome-wide background levels are shown with dashed line. c) Density heatmaps of the H3K4me1, H3K4me3, H3K9me3, H3K27ac, H3K36me3 and ATAC-signal in the 250 nt vicinity of active G4 loci in A549 cells. Top 6,000 profiles sorted by signal intensity are shown. d) Rate of decrease of epigenetic signal for the top 6,000 profiles at active G4 sites.

Recently, similar results were reported for histone mark enrichment in mouse embryonic stem cells (38), where high confidence G4 CUT&Tag peaks overlapped with open chromatin, H3K4me3 and H3K27ac peaks, whereas H3K4me1 and H3K9me3 exhibited a local minimum. Active G4, H3K4me2 and H3K4me3 peaks followed by H3K27ac and open chromatin densely occupy gene promoter regions in HEK293T cells in another study (35). It was confirmed earlier that folded G4s are enriched at gene promoter regions characterized by open chromatin (31). Therefore, H3K4me3, and H3K27ac histone modifications and open chromatin (ATAC-seq) signal are good candidates to inform the G4 prediction in cells. We aim to extract the informative signal from both the epigenetic landscape and the PQS sequence with a neural network termed *epiG4NN* and compare it with the baseline *G4NN* that only uses the DNA sequence.

### epiG4NN is a new hybrid model leveraging different epigenetic marks

We developed *epiG4NN*, a hybrid sequence and epigenetic context-based model for prediction of G4 formation in cells. *epiG4NN* is a neural network model that consists of stacks of convolutional layers with skip connections for better training convergence (58). Convolutional networks (CNNs) are a class of deep learning methods that achieved significant breakthroughs in the genomic predictions (64–67). Convolutional kernels slide along the inputs and extract input features, passed on to the next layers. CNNs allow for hierarchical representation of features through learning the patterns in the input sequence without explicit feature engineering. For inputs, we used putative quadruplex-forming sequences (PQS) and different auxiliary arrays of processed epigenetic marks aligned to PQS: histone 3 lysine 4 residue mono-methylation (H3K4me1), histone 3 lysine 4, 9, and 36 residues tri-methylation (H3K4me3, H3K9me3, H3K36me3), histone 3 lysine 27 residue acetylation (H3K27ac), and chromatin accessibility (ATAC ChIP-seq data). Each epigenetic feature was tested independently (Fig. 3). Comparisons were made with a sequence-based model *G4NN* with the same architecture and hyperparameters (number of residual stacks, convolutional kernel properties, learning rate, batch size). We trained our models on A549 G4 data from a RHAU-derived antibody G4P-ChIP-seq experiment (32) and subsequently evaluated on both A549 withheld test samples and unseen HEK293T and K562 cell data.

**Figure 3.**
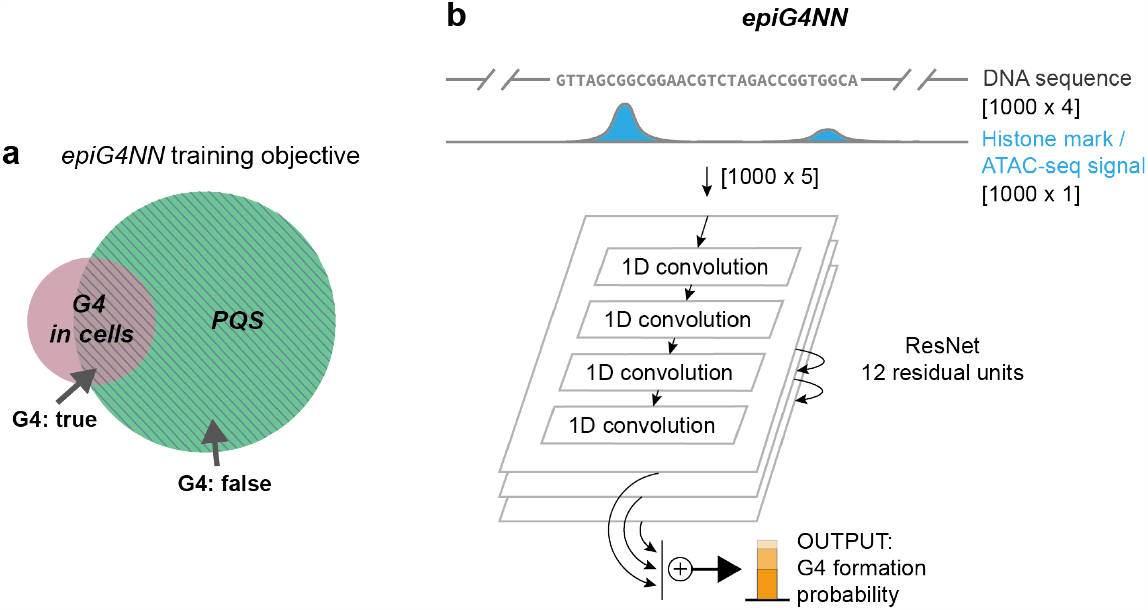
*epiG4NN* is a ResNet-based architecture directly incorporating raw DNA sequence and local epigenetic profiles for G4 prediction in cells. a) *epiG4NN* training objective is to predict putative G4 sequence (PQS) formation in cells, including those G4 not found *in vitro*. b) PQS sequence with immediate flanks to a total of 1000 nt is taken for input and stacked with the normalized array of the given epigenetic feature. The architecture is based on 12 residual convolutional blocks with dilation. *epiG4NN* outputs a normalized probability of G4 formation.

### Accurate G4 formation prediction in cells based on epigenetic features with epiG4NN

Upon optimization of hyperparameters, the best *epiG4NN* architecture was determined. We trained six *epiG4NN* models (*epiG4NN*-H3K4me1, *epiG4NN*-H3K4me3, *epiG4NN*-H3K9me3, *epiG4NN*-H3K27ac, *epiG4NN*-H3K36me3, *epiG4NN*-ATAC) on A549 cell data (32) along with the baseline sequence-only *G4NN*. Different epigenetic marks and ATAC-seq signal were used for training by independently stacking each normalized epigenetic feature array with the one-hot-encoded sequence, hence expanding the transverse dimension of the encoded sequence from 4 to 5 (Fig. 3). Both area under the receiver operating characteristic (AUROC) and area under the precision-recall curve (AUPRC) were used as quality measures of the model. AUROC measures the separability of the positive and negative classes, where a score of 1 means perfect ability of the model to distinguish between classes. However, AUROC may be skewed by an imbalance of number of samples in classes. AUPRC measures the trade-off between precision and recall, and AUPRC of 1 means high precision and high recall. The class imbalance problem exists in G4 training data. We found about 2,100,000 of potential G4 motifs, while only about 105,000 of them are formed in A549 cells. Accuracy and AUROC metrics are, therefore, not sensitive enough for training and evaluation in this problem. Instead, AUPRC can be used as characteristic for the G4 prediction objective. Previously, the maximum AUROC of 0.988 and AUPRC of 0.309 were reported for the problem of G4 prediction in cells (42). Here, we achieve an AUROC of 0.996 and an AUPRC of 0.907 (Fig. 4) for the *epiG4NN*-H3K4me3 model on the A549 unseen chromosome set. In accordance with the widely accepted fact that G4 generally colocalize with accessible chromatin regions (31), our models performance rating support the importance of open chromatin (*epiG4NN*-ATAC, AUROC = 0.993, AUPRC = 0.837), and the epigenetic mark of active enhancers, H3K27ac (*epiG4NN*-H3K27ac, AUROC = 0.991, AUPRC = 0.838) for G4 formation. *epiG4NN*-H3K4me1 gave an intermediate improvement in the prediction (AUROC = 0.984, AUPRC = 0.693), whereas *epiG4NN*-H3K9me3 and *epiG4NN*-H3K36me3 gave only moderate improvements (AUROC = 0.984, AUPRC = 0.686 and AUROC = 0.982, AUPRC = 0.673, respectively) compared to sequence-based *G4NN* (AUROC = 0.983, AUPRC=0.668). Combining two best predictive features only resulted in a marginal performance improvement (see Supplementary Note 1).

**Figure 4.**
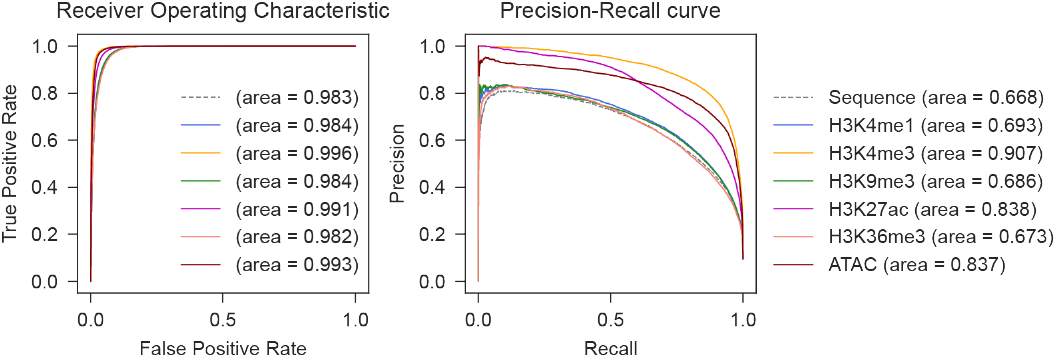
*epiG4NN* improves G4 formation prediction in cells on held-out A549 test samples by using epigenetic features for training, as measured by receiver operating characteristic and precision-recall curves. Receiver operating characteristic (left) and precision-recall (right) curves of the baseline *G4NN* model (dashed gray) and *epiG4NN*-H3K4me1, *epiG4NN*-H3K4me3, *epiG4NN*-H3K9me3, *epiG4NN*-H3K27ac, *epiG4NN*-H3K36me3, *epiG4NN*-ATAC models. Areas under the receiver operating characteristic curve (AUROC) and under the precision-recall curve (AUPRC) are shown in the legend.

### epiG4NN predicts G4 formation in unseen cell lines

Populations of G4s in cellular context are shared between cell lines or formed uniquely in some cells. We compared the pre-processed G4 peaks obtained from HEK293T, HaCaT, HeLa cells using G4P ChIP-seq (32), K562 cells using BG4 ChIP-seq (37), and K562 cells using CUT&Tag (34) with G4 peaks detected in A549 cells with G4P ChIP-seq (32). A549 and HeLa cell lines are epithelial-like cells derived from lung and uterus, respectively, whereas HaCaT are keratinocytes; HEK293T are derived from kidney cells, and K562 are lymphoblast cells. G4P ChIP-seq and CUT&Tag methods detect G4s *in situ*, while BG4 ChIP-seq detects G4s *ex vivo*. We found that only 53% of HaCaT, 76% of HEK293T, 60% of K562 (BG4 ChIP-seq), 38% of K562 (CUT&Tag) and 82% of HeLa G4 peaks are common with A549. To assess how well our model performs on different cell types with distinct underlying epigenetic landscapes, we additionally carried out model evaluation on HEK293T (G4P ChIP-seq) and K562 (CUT&Tag) cell lines. As *epiG4NN* was trained on G4 peaks from A549 cells obtained with G4P ChIP-seq, we only used data from *in situ* G4 detection experiments to exclude possible technical variability. We selected the three best models as tested on A549 unseen samples (*epiG4NN*-H3K4me3, *epiG4NN*-ATAC, *epiG4NN*-H3K27ac) together with the baseline sequence-only model *G4NN* and measured their performance on new cell lines with AUROC and AUPRC (Fig. 5). *epiG4NN*-H3K4me3 showed the best performance with both HEK293T (AUPRC = 0.836) and K562 (CUT&Tag) (AUPRC = 0.838) cell data, followed by *epiG4NN*-ATAC for K562 (CUT&Tag) and *epiG4NN*-H3K27ac for HEK293T. *epiG4NN*-ATAC, however, performed slightly worse than sequence alone for HEK293T.

**Figure 5.**
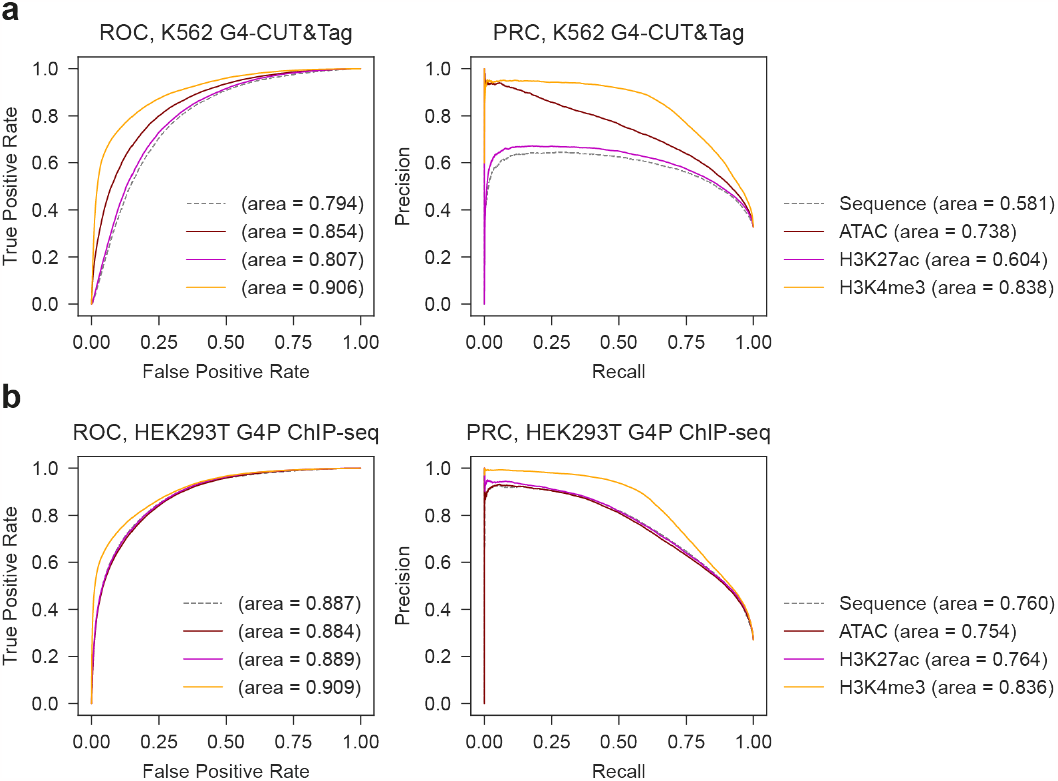
*epiG4NN* predicts G4 formation in unseen cell lines. Receiver operating characteristics and precision-recall curves for *epiG4NN* evaluation of G4 formation prediction: a) in K562 cells obtained with an *in situ* CUT&Tag experiment (34), b) in HEK293T cells obtained in an *in situ* G4P ChIP-seq experiment (32), all evaluated with *G4NN* (sequence-only, gray dashed line) and *epiG4NN*-ATAC, *epiG4NN*-H3K27ac, and *epiG4NN*-H3K4me3. Areas under the receiver operating characteristic curve (AUROC) and under the precision-recall curve (AUPRC) are shown in the legend.

### Model learned from in situ data cannot predict ex vivo data well

Additionally, we tested another K562 cell line G4 dataset obtained with the *ex vivo* BG4 ChIP-seq method. Given the same epigenetic profile and the same sequence motifs as for the K562 (CUT&Tag) data, with a different set of G4 peaks to predict, *epiG4NN*-H3K4me3 and *epiG4NN*-H3K27ac models were only able to perform marginally better than the *G4NN* baseline in terms of AUROC and AUPRC; *epiG4NN*-ATAC resulted in a slightly better evaluation (AUROC = 0.909, AUPRC = 0.342) (Fig. 6). The underlying technical/experimental condition difference between BG4 ChIP-seq and CUT&Tag methods of G4 detection in cells, therefore, makes the transfer of learned features challenging. The features learnt from one experimental condition may differ from the features in other conditions, resulting in poor cross-method model performance.

**Figure 6.**
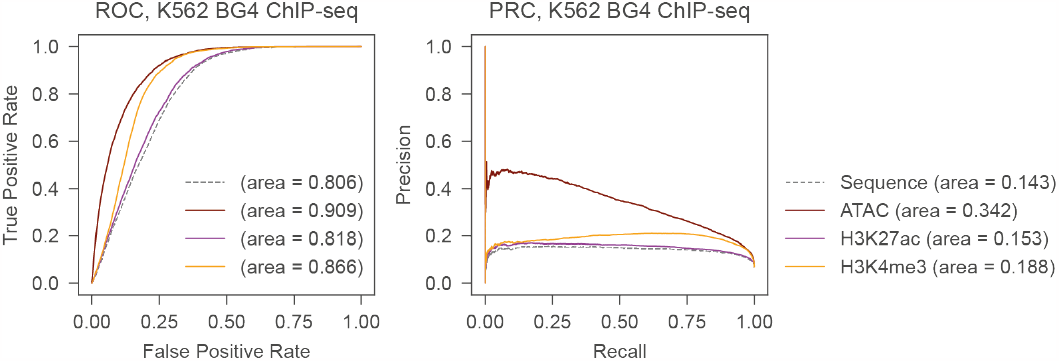
*epiG4NN* has compromised performance in an ex vivo G4 dataset. *epiG4NN*-ATAC, *epiG4NN*-H3K27ac, and *epiG4NN*-H3K4me3 models were evaluated on BG4 ChIP-seq data of K562 cells with receiver operating characteristics and precision-recall curves. Areas under the receiver operating characteristic curve (AUROC) and under the precision-recall curve (AUPRC) are shown in the legend.

### An example of differential epiG4NN prediction

The first intronic region of the human *GRHL3* gene, located on the chromosome 1, contains a few PQS, where one PQS is formed in both HEK293T and A549 cell lines, while the other PQS is formed in HEK293T cells but not in A549 cells (32) (Fig. 7). We created pseudo-genomic tracks that demonstrate the formation of these PQS using *epiG4NN*-H3K4me3 and making point predictions for each nucleotide in the region of interest.

**Figure 7.**
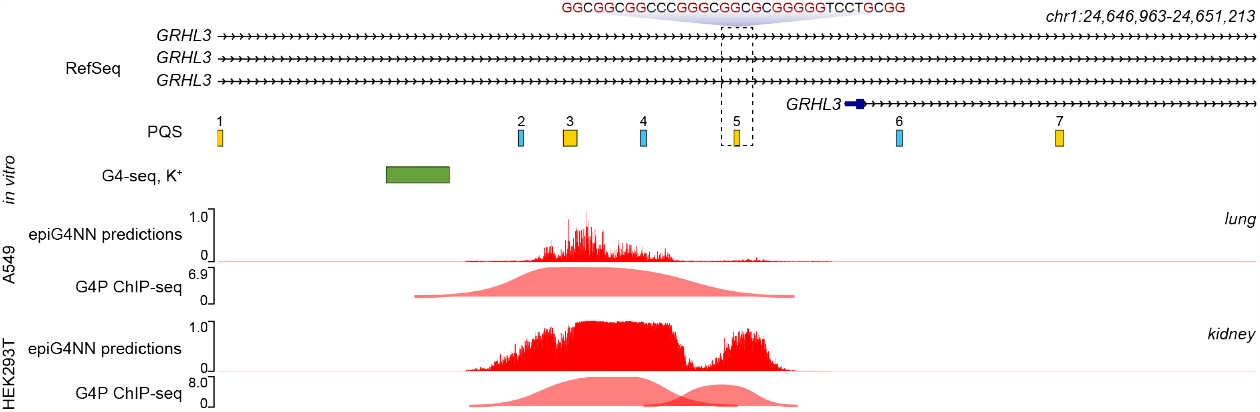
*epiG4NN* predicts universally and differentially formed G4 in the *GRHL3* first intron region. Genome tracks, from top to bottom: 1) RefSeq transcripts; PQS detected in *hg19* with bioinformatic motif search, plus strand: yellow, minus strand: blue; 2) *in vitro* detected G4 peaks (G4-seq in K^+^) (31), plus strand: green, minus strand: not detected; 3) A549 predictions with *epiG4NN*-H3K4me3 and G4 peaks detected with G4P ChIP-seq (32); 4) HEK293T predictions with *epiG4NN*-H3K4me3 and G4 peaks detected with G4P ChIP-seq (32). PQS numbered 3 has a peak in both HEK293T and A549 cells, while PQS number 5 has a peak in HEK293T only. PQS number 5 is highlighted, and its sequence is shown.

We found seven PQS corresponding to *GRHL3* intron 1 region (Fig. 7), where four PQS are found on the plus strand (numbered 1, 3, 5, and 7), and three PQS on the minus strand (numbered 2, 4, and 6). We additionally refer to the *in vitro* data. Interestingly, only one *in vitro* formed G4 on the plus strand was matching the region, and it did not overlap with any of the PQS motifs. In cells, A549 G4P ChIP-seq data has one G4 peak corresponding to PQS number 3 and HEK293T has two peaks corresponding to PQS numbers 3, 5 (Fig. 7). *epiG4NN*-H3K4me3 captures the difference between the H3K4me3 features in A549 and HEK293T cell lines given the same sequence of the G4 motif and predicts unique formation of the PQS number 5 in HEK293T, while PQS number 3 is predicted in both cell lines. Previously reported DeepG4 model only used G4 forming both in cells and *in vitro* for training and testing (42). The lack of matching *in vitro* peak highlights the importance of prediction in cells irrespective of the G4 formation *in vitro*.

### epiG4NN-H3K4me3 exhibits a promoter and enhancer bias

Certain histone marks are known to be enriched in open chromatin, gene enhancer, and promoter regions of the genome (68). H3K4me3 is an “active” histone mark thought to play a role in transcription (69, 70) and is marking gene promoters (71), while H3K27ac is marking gene enhancers (72). To test whether our model is biased towards these regions, we extracted G4 overlapping gene promoters (73) and cell-specific enhancers for A549 and HEK293T cell lines (74), and evaluated the region-specific performance of *epiG4NN*-H3K4me3. Evaluation revealed overall better AUPRC scores for enhancer and promoter regions compared to random regions (Fig. 8) together with the fact that a greater proportion of G4 is formed in promoter regions. Ubiquitous formation of G4 in the promoter regions was indeed experimentally confirmed previously (31, 54, 75). A significantly lower proportion of the G4 is formed in the enhancer regions (Fig. 8d), and even less so in the random regions (Fig. 8f), while the quality of prediction drops the most drastically for random regions in HEK293T cell lines.

**Figure 8.**
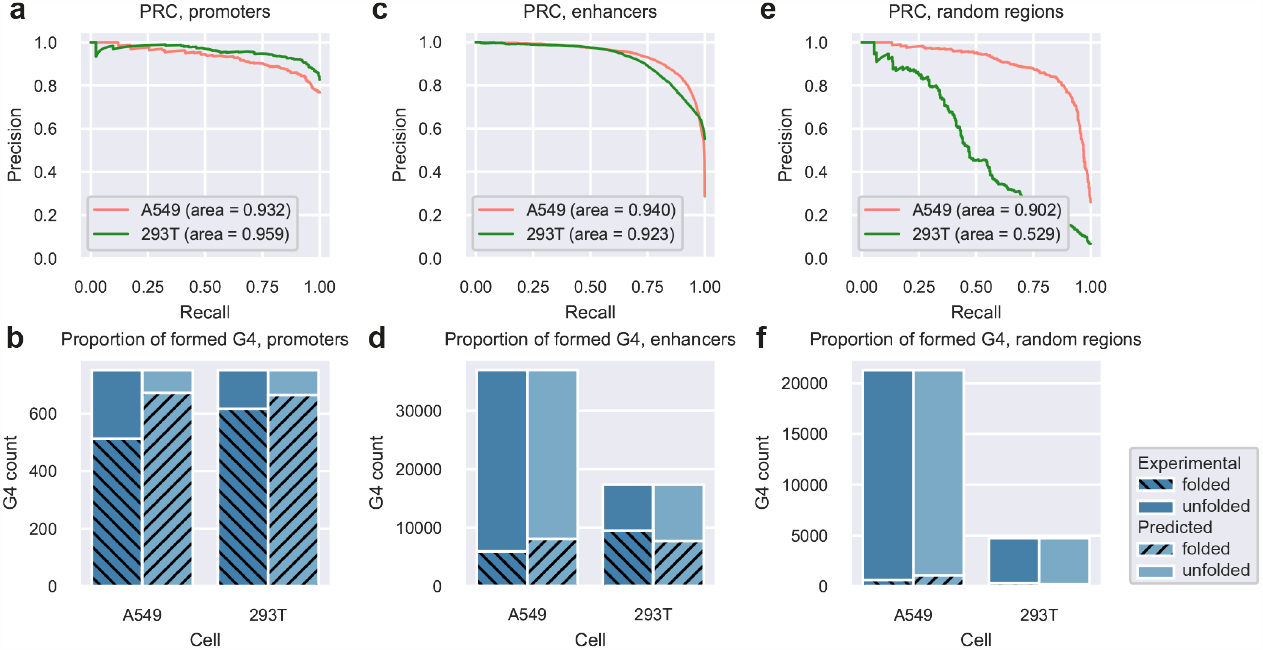
Promoter-enhancer bias in the *epiG4NN* prediction quality. a), c), e) precision-recall curves for *epiG4NN*-H3K4me3 in promoters, enhancers, and random regions in A549 (left-out chromosome set) and HEK293T (unseen) cells, respectively. b), d), f) proportions of formed G4, experimental and predicted, in promoters, enhancers, and random regions in A549 and HEK293T cells.

## DISCUSSION

Genome-wide G4 prediction methods are important for understanding G4 biology and for targeting such structures. Recent progress in chromatin immunoprecipitation methods was applied to G4 detection, and multiple experimental G4 datasets were reported. However, the discrepancies between G4 formed in different cells and experiment types pointed to the need for understanding G4 formation in cells. The vast body of cellular epigenetic data allows to attribute G4 formation in cells to specific cellular features. So far, little is known about the correlations between G4 formation and such cellular data. Here, we demonstrate a novel approach, *epiG4NN*, that comprises of a hybrid deep neural network that uses cellular epigenetic features and DNA sequence for G4 prediction in genomic DNA. Compared to previously published methods, *epiG4NN* achieves unprecedented precision and recall in G4 prediction in unseen cell lines. Additionally, *epiG4NN* allows to rate the relevance of H3K4me1, H3K4me3, H3K9me3, H3K27ac, H3K36me3 and chromatin accessibility signals to G4 prediction problem.

Through our experiments on the A549, HEK293T and K562 data, we show that supplementing epigenetic data improves learning, as compared to sequence-based model *G4NN* with the same architecture and number of parameters. We demonstrate that *epiG4NN*-H3K4me3, *epiG4NN*-H3K27ac and *epiG4NN*-ATAC considerably outperform *G4NN*. H3K4me3 is the strongest predictor of the G4 formation in our experiments on A549 and HEK293T (G4P ChIP-seq (32)) and K562 cell data (CUT&Tag (34)), followed by ATAC-seq and H3K27ac signals. We found that the optimal epigenetic predictor for cell lines depends on the experimental condition of the underlying data. Independent evaluation of *epiG4NN* on G4 data obtained with G4P ChIP-seq (32) and CUT&Tag (34) showed that H3K4me3 is a good predictor of G4 formation for the G4 detected *in situ*, while open chromatin is the only predictor for *ex vivo* type of experiment (K562 data, BG4 ChIP-seq (37)) that improves over the sequence-only prediction. This likely reflects the different approach behind the experimental datasets: K562 cells in BG4 ChIP-seq were fixed and chromatin was fragmented before it reacted with a G4 specific antibody, while A549 cells were subjected to a G4P knock-in and the antibody was expressed natively. It is not clear how fragmentation and purification of DNA affects G4 formation, and features learned by the model trained on *in situ* data do not seem to translate to another class of G4 detection experiment. The finding that ATAC-seq improves G4 predictions efficiency is in line with a previous report (42), while H3K4me3 was found to be highly colocalized with G4 sites in other recent studies (35, 38). Additionally, we have demonstrated that only H3K4me3, H3K27ac and open chromatin signals contribute to a large number of active G4 sites, while H3K4me1, H3K9me3 and H3K36me3 are largely depleted. The key difference between *epiG4NN* and previously reported models lies in the usage and comparison of multiple epigenetic marks or features for contextual prediction of G4 formation in cells. We achieve a better performance in G4 prediction and demonstrate relative importance of different epigenetic features. We additionally retrained the previously reported model DeepG4 (42) on our data to compare the model architectures, and obtained an AUROC or 0.981 and an AUPRC of 0.644 for the A549 left-out test samples (Supplementary Fig. 5), demonstrating that our architecture may be more suitable for this task. Additionally, unlike DeepG4 – the only other model predicting DNA G4 in cells reported so far – we employ a full snapshot of the local epigenetic profile, in contrast with a single average value for a given G4 motif region. We show that our model can reproduce peak signatures from two different cell lines, A549 and HEK293T, where a PQS is formed differentially. Our model, however, suffers from a prediction accuracy bias in the random regions as compared with gene promoter or enhancer regions. We believe that *epiG4NN* can contribute to study the roles of chromatin marks on the sequence-structure dependence.

## DATA AVAILABILITY

Data preparation and *epiG4NN* training scripts are available in the GitHub repository (https://github.com/anyakors/epiG4NN).

## SUPPLEMENTARY DATA

Supplementary Data are available at NAR online.

## ACKNOWLEDGEMENT

The authors thank Jasraj Singh for his help in preparing the scripts for epigenetic data pre-processing.

## FUNDING

Nanyang Technological University (NTU Singapore) grants (to A.T.P.). Funding for open access charge: Nanyang Technological University.

## CONFLICT OF INTEREST

The authors have no competing interests to declare.

## Notes

### Competing Interest Statement

The authors have declared no competing interest.

