## Supplementary Data for "Prediction of G4 formation in live cells with epigenetic data: a deep learning approach"

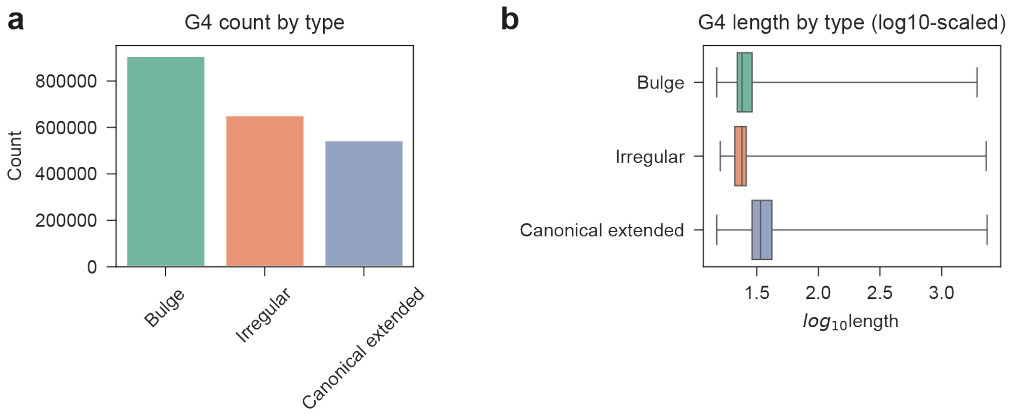


Supplementary Figure 1: Bioinformatic G4 search in the hg19 human reference genome revealed N=2,105,837 G-quadruplexes. a) Count of G4 by type: bulged, irregular and canonical extended G4. b) Boxplot of $\log_{10}$ of the G4 length by type.

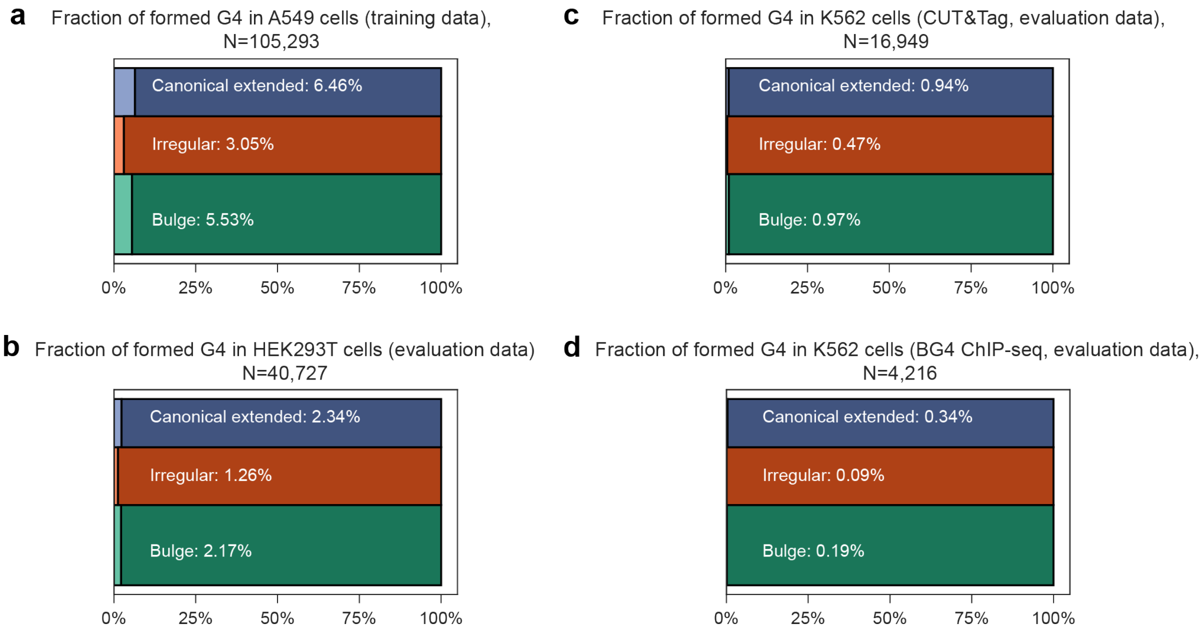


Supplementary Figure 2: Fractions of formed G4 (positive label) and G4 that are not formed (negative label) by type: a) A549 cells (training data), b) HEK293T cells (evaluation data), c) K562 cells, CUT&Tag (evaluation data), c) K562 cell, BG4 ChIP-seq (evaluation data).

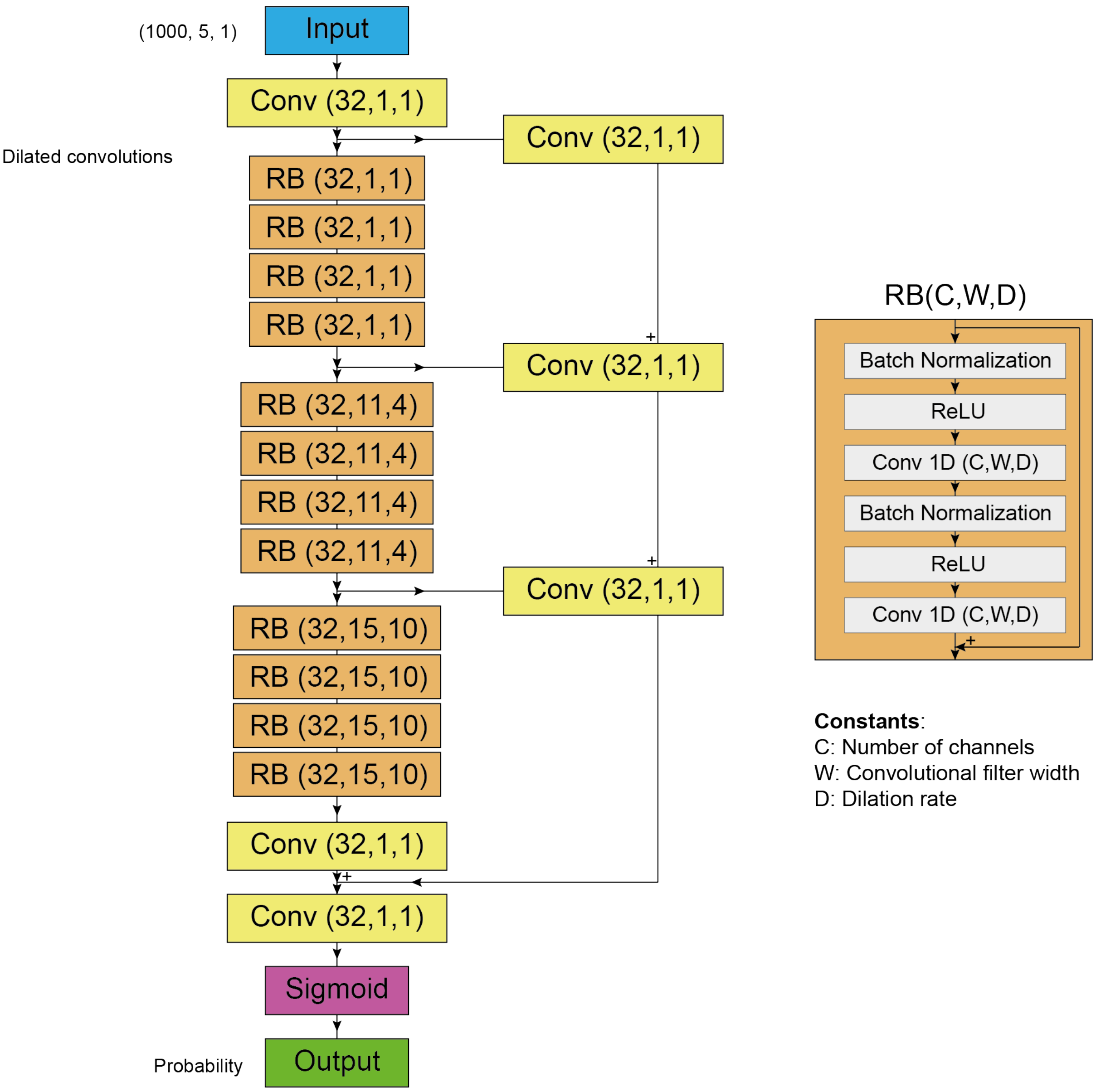


Supplementary Figure 3: *epiG4NN* architecture is a deep convolutional neural network with dilation. *epiG4NN* consists of three towers of four residual blocks each. The layers of the residual block are shown on the right. Batch state is saved before each convolutional tower and added before the penultimate convolution.

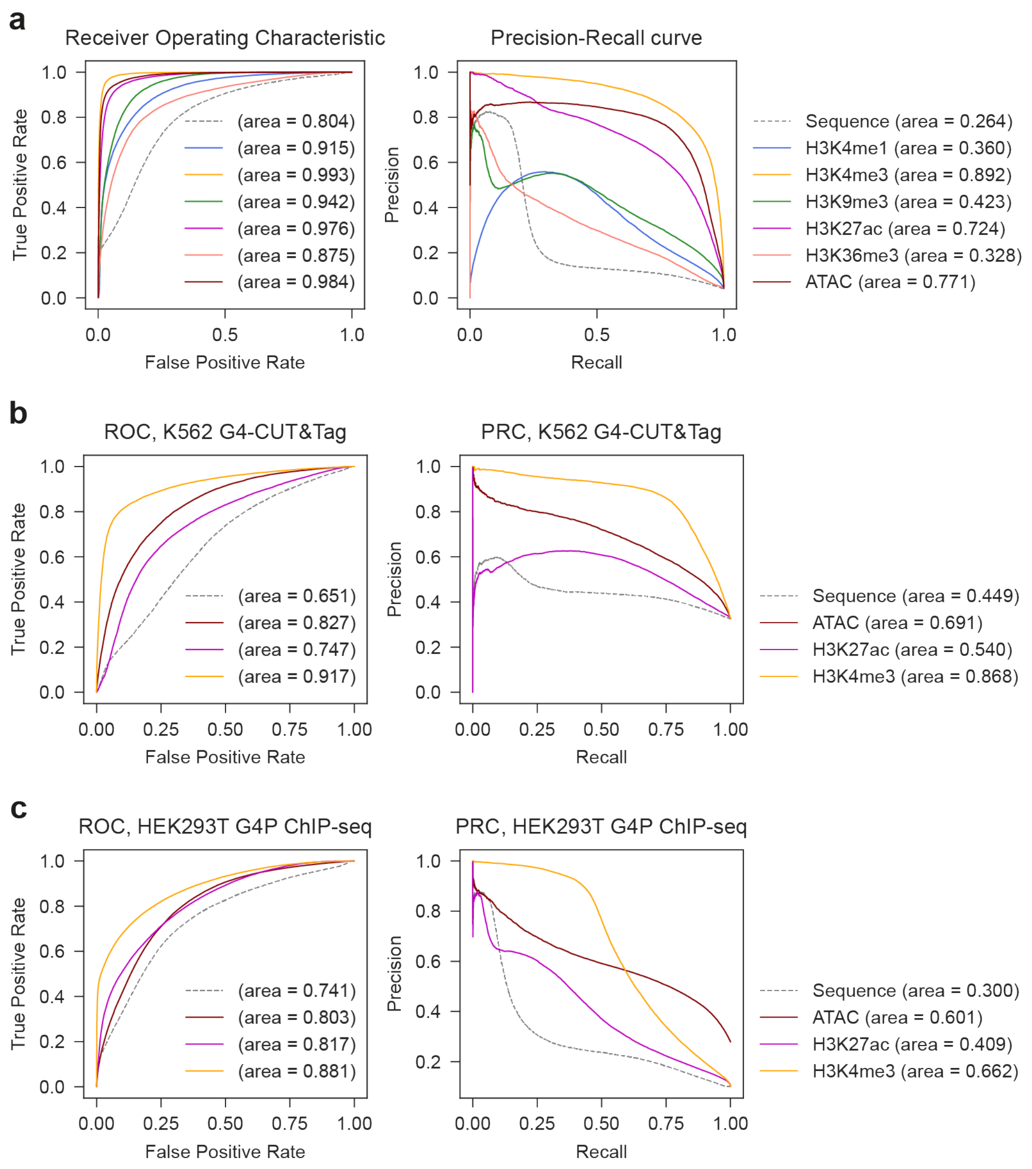


Supplementary Figure 4: *epiG4NN*-250nt performance on the test set from the cell line used for training (A549 (a)) and on independent data (K562 (b), HEK293T (c)). Instead of 1000nt stretches of sequences and respective epigenetic data, 250nt-long inputs were used.

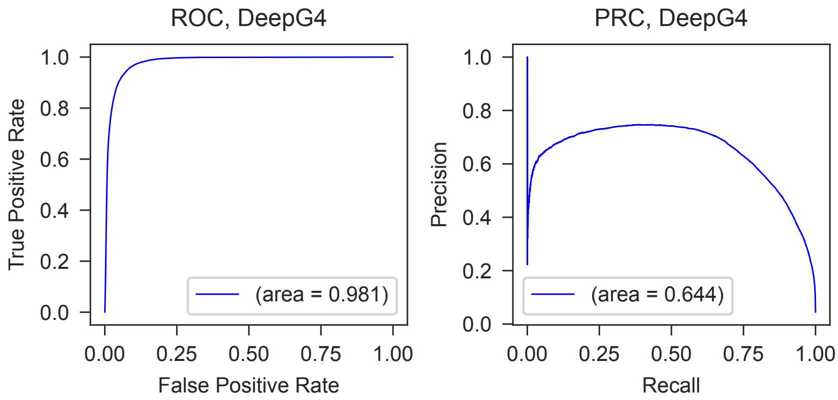


Supplementary Figure 5: We retrained the DeepG4 model (1) with the reported architecture on the pre-processed A549 data, and obtained performance metrics higher than reported originally in (1).

### Supplementary Note 1

***Combination of chromatin accessibility and H3K4me3 inputs does not improve the performance significantly***. We combined H3K4me3 and ATAC-seq signal for the 250nt model input to test if combining the best performing additional layers will result in a synergetic model. The resulting model's performance, however, was only marginally better on both A549 left-out test samples (AUROC = 0.995, AUPRC = 0.897), as compared to the *epiG4NN*-H3K4me3 model alone (AUPRC = 0.892, see Supplementary Fig. 4). This suggests a potential improvement in the epigenetic feature extractor for the architecture, as simply layering the epigenetic profiles may not result in the latent variable extraction, and a separate epigenetic input processing head may be needed.
